# A mouse model of occult intestinal colonization demonstrating antibiotic-induced outgrowth of carbapenem-resistant *Enterobacteriaceae*

**DOI:** 10.1101/2021.02.24.432587

**Authors:** Choon K. Sim, Sara Saheb Kashaf, Sean Conlan, Apollo Stacy, Diana M. Proctor, Alexandre Almeida, Nicolas Bouladoux, Mark Chen, NISC Comparative Sequencing Program, Robert D. Finn, Yasmine Belkaid, Julia A. Segre

## Abstract

**Background:** The human intestinal microbiome is a complex community that contributes to host health and disease. In addition to normal microbiota, pathogens like carbapenem-resistant *Enterobacteriaceae* may be asymptomatically present. When these bacteria are present at very low levels, they are often undetectable in hospital surveillance cultures, known as occult or subclinical colonization. Through the use of antibiotics, these subclinical pathogens can increase to sufficiently high levels to become detectable, in a process called outgrowth. However, little is known about the interaction between gut microbiota and *Enterobacteriaceae* during occult colonization and outgrowth.

**Results:** We developed a clinically relevant mouse model for studying occult colonization. Conventional wild-type mice without antibiotic pre-treatment were exposed to *K. pneumoniae* but rapidly tested negative for colonization. This occult colonization was found to perturb the microbiome as detected by both 16S rRNA amplicon and shotgun metagenomic sequencing. Outgrowth of occult *K. pneumoniae* was induced either by a four-antibiotic cocktail or by individual receipt of ampicillin, vancomycin or azithromycin, which all reduced overall microbial diversity. Notably, vancomycin was shown to trigger *K. pneumoniae* outgrowth in only a subset of exposed animals (outgrowth-susceptible). To identify factors that underlie outgrowth susceptibility, we analyzed microbiome-encoded gene functions and were able to classify outgrowth-susceptible microbiomes using pathways associated with mRNA stability. Lastly, an evolutionary approach illuminated the importance of xylose metabolism in *K. pneumoniae* colonization, supporting xylose abundance as a second susceptibility indicator. We showed that our model is generalizable to other pathogens, including carbapenem-resistant *Escherichia coli* and *Enterobacter cloacae*.

**Conclusions:** This study suggests that microbiota mRNA and small-molecule metabolites may be used to predict outgrowth-susceptibility. Our modeling of occult colonization and outgrowth could help the development of strategies to mitigate the risk of subsequent infection and transmission in medical facilities and the wider community.

## Introduction

It is well-established that the gut microbiome plays a critical role in the health of the host [1, 2]. Conditions like irritable bowel syndrome and Crohn’s disease have been linked to dysbiosis of the microbiome [3], as have other complex conditions like obesity and diabetes [4]. The commensal microbiota also plays a critical role in ‘colonization resistance,’ or the prevention of pathogenic organisms from taking up residence in the gut, from which they could spread to other organs [5, 6]. Factors that disrupt the microbiome, such as antibiotics [7], nutritional changes [8] or competition between microbes [9], can alter the microbiome’s population structure, which may allow for colonization or outgrowth by pathogenic organisms.

Outgrowth of pathogenic bacteria, like carbapenem-resistant *Enterobacteriaceae* (CRE), is of particular concern in hospitals and other healthcare settings. CRE such as *Klebsiella pneumoniae, Escherichia coli* and *Enterobacter cloacae* are an important cause of antibiotic-resistant infections worldwide [10]. According to a 2019 report from the US Centers for Disease Control and Prevention, CRE are considered an urgent threat, resulting in 13,100 hospitalizations, 1,100 deaths, and an estimated $130 million in healthcare costs in the US annually [11]. This has led hospitals to implement intake screening protocols to limit the spread of CRE when patients are admitted or transferred from other facilities [12, 13]. Screening allows patients with active infection or asymptomatic colonization [14, 15] to be detected and properly isolated from vulnerable populations.

Confounding screening efforts, patients with low-level, sub-clinical (occult) colonization are not easily detected by intake screenings or routine surveillance cultures [16]. Outgrowth from occult colonization is hard to differentiate from nosocomial transmission without detailed molecular analyses [17, 18]. Robust methods to distinguish between these two scenarios would have important clinical implications. While nosocomial transmission would warrant extensive screening of an entire hospital unit, CRE outgrowth in a single patient could be addressed as an isolated event.

Occult colonization can also play a role in community spread of CRE. CRE can persist in the gastrointestinal tract of healthy individuals and the general population [19]. Travelers to CRE-endemic regions are at risk of being exposed to CRE, even without visiting a medical facility, and they may carry it back to their home country upon return from travel [20, 21].

Occult colonization is, by definition, difficult to detect directly, but we hypothesized that there might be other signals that indirectly report on low-level colonization and subsequent risk of outgrowth. In the current study, we constructed a mouse model for occult colonization and used both 16S rRNA and shotgun metagenomic sequencing to demonstrate that occult colonization remodels the microbiome. We then identified vancomycin receipt as a condition under which we could achieve incomplete outgrowth after antibiotic treatment from an undetectable state. This allowed us to identify functions involved in mRNA stability that were enriched in mice where outgrowth was observed. Finally, we identified mutations that evolved in *K. pneumoniae* under vancomycin selection, suggesting xylose may play a role in outgrowth.

## Results

### Development of a murine model of occult colonization

We established a murine model to identify markers of occult intestinal colonization by multidrug-resistant *Enterobacteriaceae*. We chose C57BL/6N wild-type mice and a dominant lineage, sequence type 258 (ST258) strain of *K. pneumoniae* (KPNIH1) [18, 22] for this model. Furthermore, ST258 has been demonstrated to colonize patients long-term [23], and it has a well-curated transposon library of knockout mutants [24]. We initially studied 3 variables: inoculum size, location of occult colonization and time until antibiotics (**Fig. 1a**). Most infection models use a very large inoculum (>10^7^ CFU), but as we were interested in occult colonization, we chose a more conservative range of inoculum sizes. We titrated from 10^3^ to 10^5^ CFU and observed at days 4-6 post-gavage that 10^5^ CFU inoculum was rarely detectable, whereas the 10^4^ and 10^3^ CFU inocula were never detectable in stool (**Fig. 1b**). To determine if mice were colonized at levels below our detection threshold, we administered antibiotics, as this has been shown to trigger outgrowth of resistant strains [25]. Initially, we chose an antibiotic cocktail, comprising ampicillin, vancomycin, neomycin and metronidazole. The KPNIH1 strain we chose to work with is resistant to all 4 of these antibiotics (**Additional file 1a**), and this cocktail is widely used in murine studies to ablate microbiota [26]. Administering the antibiotic cocktail 1 week post-gavage allowed *K. pneumoniae* to grow from an occult state to high levels in stool (**Fig. 1b**). Of note, because we chose to singly house mice, *K. pneumoniae* outgrowth was not a result of mouse-to-mouse transmission.

**Fig. 1.**
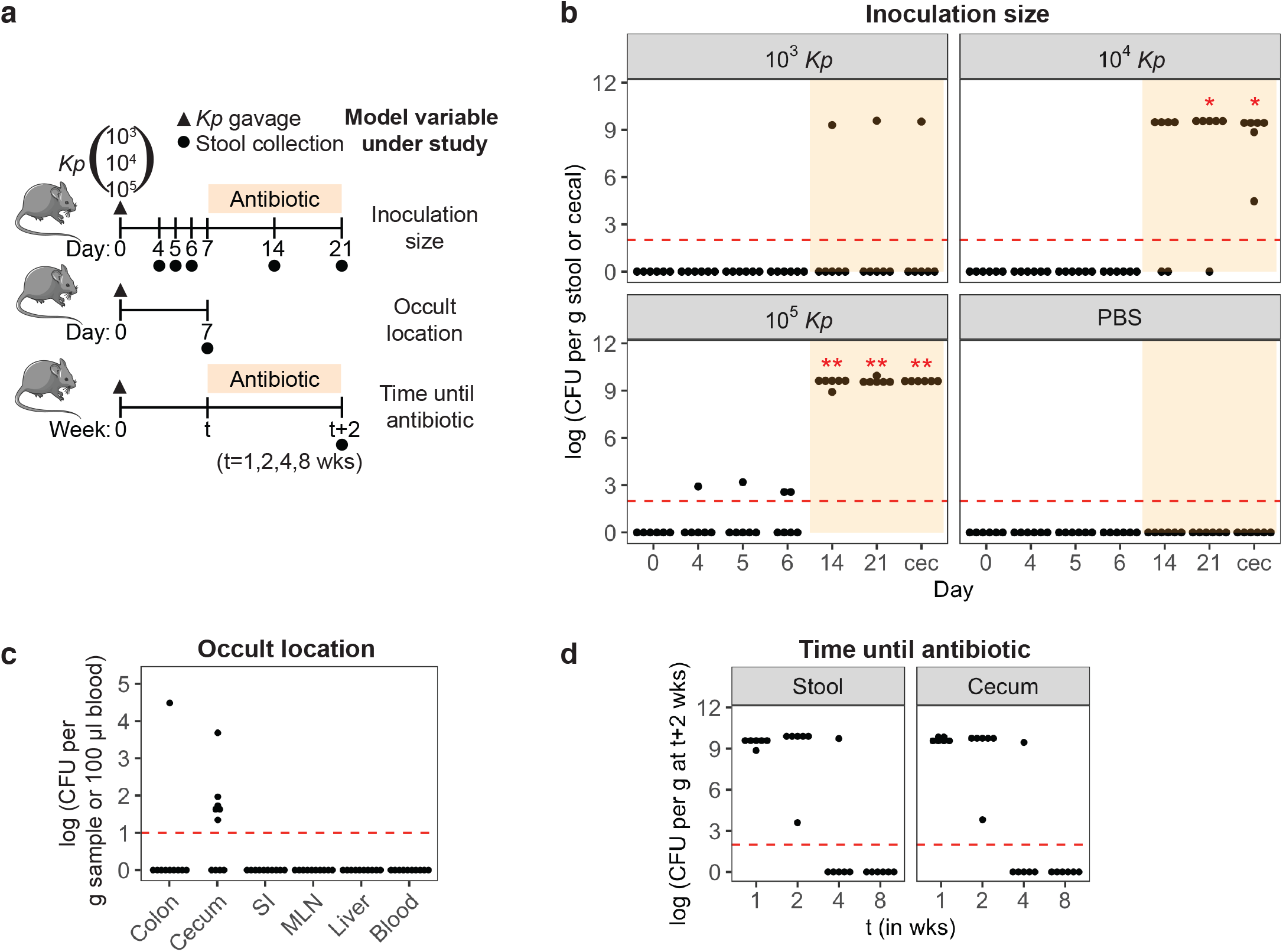
*K. pneumoniae* occultly colonizes the gastrointestinal tract for up to 4 weeks, with outgrowth following receipt of an antibiotic cocktail. **a** Study designs to test inoculation size, occult location and time until antibiotic. (Top) Mice were orally gavaged with 10^3^, 10^4^, or 10^5^ CFU of *K. pneumoniae* (*Kp*) or the PBS vehicle control at day 0 (triangle). An antibiotic cocktail was given from day 7 to day 21. Days of stool collection are marked by circles. Mice were euthanized at day 21, and cecal contents collected. (Middle) Mice were orally gavaged with 10^5^ CFU of *K. pneumoniae*, and tissues were plated at day 7. (Bottom) Mice were orally gavaged with 10^5^ CFU of *K. pneumoniae*, and different groups of mice were given antibiotic for 2 weeks starting at t = 1, 2, 4 or 8 weeks post-gavage. Stool and cecal contents were collected at (t+2)^th^ week. **b** CFU of *K. pneumoniae* in stool and cecal contents (cec). N=6. The red dotted line marks the limit of detection for CFU. **c** CFU of *K. pneumoniae* in colonic, cecal and small intestinal (SI) contents and the mesenteric lymph nodes (MLN), liver and blood plasma. N=10. **d** CFU of *K. pneumoniae* at (t+2)^th^ weeks in stool and cecal content. N=6. For (**b**), Fisher’s exact test with Hommel correction for multiple testing was used to compare numbers of mice showing outgrowth versus no-outgrowth in inoculated mice to those in the PBS control group. Outgrowth is defined as CFU > 0. \**p*-value < 0.05, \*\**p*-value < 0.01

Next, we determined the lowest inoculation dose possible for *K. pneumoniae* outgrowth from an occult state. While outgrowth was observed for the 10^5^ CFU inoculum in all mice, 10^4^ CFU resulted in outgrowth in most but not all mice. 10^3^ CFU very rarely resulted in outgrowth, suggesting that *K. pneumoniae* is typically unable to colonize occultly from such a starting inoculum (**Fig. 1b**). Thus, we chose 10^5^ CFU as the inoculum for subsequent experiments.

*K. pneumoniae* is capable of colonizing many body sites, and some strains can translocate across the gut barrier [27]. To identify where in the host *K. pneumoniae* establishes occult colonization, we examined the gastrointestinal tract as well as peripheral tissues at day 7 post-gavage (**Fig. 1a**, middle). We identified *K. pneumoniae* in the cecum of the majority of mice, but not the small intestines, mesenteric lymph nodes, liver or blood plasma (**Fig. 1c**), suggesting that the primary reservoir of *K. pneumoniae* occult colonization is the cecum with no systemic spread.

To determine the duration of *K. pneumoniae* occult colonization, we initiated receipt of antibiotics at 1, 2, 4 or 8 weeks post-inoculation (**Fig. 1a**, bottom). We observed that *K. pneumoniae* consistently showed outgrowth when antibiotic was administered at 1 or 2 weeks but rarely if at all at 4 or 8 weeks, suggesting that, in our model, *K. pneumoniae* occultly colonizes the gut for up to 4 weeks (**Fig. 1d**). Our optimized occult colonization model using an inoculation dose of 10^5^ CFU and administration of antibiotic at 1 week post-gavage for 2 weeks duration (**Fig. 2a**), mimics a clinically relevant timecourse of bacterial outgrowth. Having a robust model is our first step to finding markers for occult colonization and outgrowth.

**Fig. 2.**
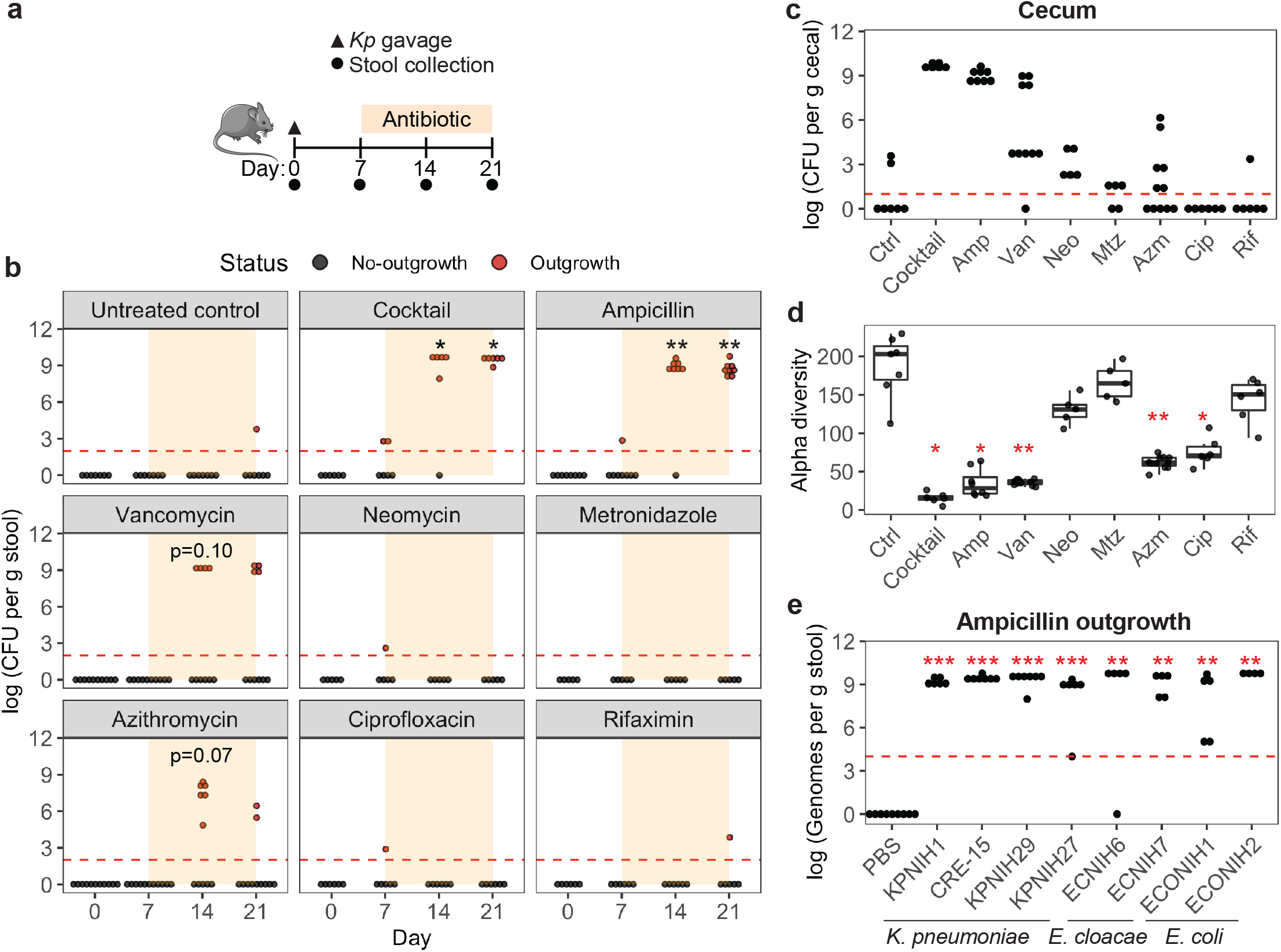
Ampicillin, vancomycin and azithromycin induce outgrowth of occult *K. pneumoniae*. **a** Mice were orally gavaged with 10^5^ CFU of *K. pneumoniae* at day 0 (triangle) and treated with antibiotic at day 7 for 2 weeks (orange region). Days of stool collection are marked by circles. **b**, **c** CFU of *K. pneumoniae* in (**b**) stool at days 0, 7, 14, and 21 and (**c**) cecal contents at day 21. The red dotted line marks the limit of detection for CFU. N=5-11. **d** Alpha diversity (number of amplicon sequence variants or ASVs) of antibiotic-treated microbiota at day 14. **e** Same experiment as (**a**), but the bacteria gavaged were other carbapenemase producing strains of *K. pneumoniae*, *E. cloacae* or *E. coli*, followed by ampicillin treatment. Levels of *Enterobacteriaceae* in stool at day 21. Measurement was done by qPCR using primers at *bla*_KPC_ gene and Ct values were converted to number of genomes using a standard curve made from dilutions of known DNA quantity. N=4-9. For (**b**) and (**e**), Fisher’s exact test with Hommel correction for multiple testing was used to compare numbers of mice showing outgrowth versus no-outgrowth in inoculated mice to those in the PBS control group. Outgrowth is defined by detected CFU > 0. For (**d**), Kruskal-Wallis H test was performed to test significance, followed by pairwise Bonferroni-corrected Mann-Whitney U tests to compare each treatment group against the control group. \**p*-value < 0.05, \*\**p*-value < 0.01, \*\*\**p*-value < 0.001.

### Clinically relevant antibiotics ampicillin, vancomycin and azithromycin induce outgrowth of *K. pneumoniae*

We reasoned that determining which individual antibiotic(s) in the 4-drug cocktail is responsible for driving outgrowth could provide insight into the underlying biology. To this end, we systematically tested each of the antibiotics in the cocktail for whether they could individually induce outgrowth (**Additional file 1b**). Among these, ampicillin and vancomycin are known clinically to predispose individuals to *K. pneumoniae* colonization [28, 29]. Accordingly, we observed significant outgrowth of *K. pneumoniae* after treatment with ampicillin (*p* < 0.01), a trend towards outgrowth after vancomycin (*p =* 0.10 at day 14), but no outgrowth with metronidazole or neomycin. We also tested the clinically relevant antibiotics ciprofloxacin, rifaximin and azithromycin (**Additional file 1b**) since these drugs are commonly used by travelers in CRE-endemic regions [30]. Among these, only azithromycin produced a trend towards *K. pneumoniae* outgrowth (*p* =0.075 at day 14) (**Fig. 2b**). Importantly, the resistance of *K. pneumoniae* to all the antibiotics tested (**Additional file 1a**) and the retention of *K. pneumoniae* in the cecum of many no-outgrowth mice (**Fig. 2c**) ruled out the possibility that *K. pneumoniae* was killed by the antibiotics or that occult colonization failed. More likely, outgrowth is influenced by antibiotic-specific perturbations to the microbial community in the gut.

To probe the general effect of 1 week of antibiotics on the microbiota, we used 16S rRNA sequencing to measure microbial alpha diversity (measure of the ecological richness within a single sample). In agreement with past studies [7, 26, 31, 32], low alpha diversity was associated with the antibiotic cocktail as well as individual ampicillin, vancomycin and azithromycin treatments (*p* < 0.05; **Fig. 2d**). Together, these results suggest that antibiotics that disrupt microbial community structure tend to promote *K. pneumoniae* outgrowth.

Next, to assess whether antibiotic outgrowth is a general feature of *Enterobacteriaceae* occult colonization, we tested whether ampicillin treatment would result in outgrowth from an occult state by other carbapenemase producing strains of *K. pneumoniae* as well as carbapenemase producing strains of *Enterobacter cloacae* and *Escherichia coli* [17, 33, 34] (**Fig. 2e**). We observed significant outgrowth of all strains tested upon ampicillin receipt (*p* < 0.01 at day 21), demonstrating that occult colonization (**Additional file 2**) and antibiotic-induced outgrowth (**Fig. 2e**) are generalizable to the *Enterobacteriaceae* family.

### Occult colonization with *K. pneumoniae* perturbs the structure of the microbiome

We hypothesized that the occultly colonized state could be accompanied by a shift in other members of the microbial community. We used 16S rRNA sequencing to compare baseline (day 0) stool samples with stool from mice occultly colonized with *K. pneumoniae* at day 7. We observed a significant reduction in the abundance of the *Erysipelotrichaceae* family (*p* = 8 x 10^−8^; **Additional file 3a**) and its component genus *Dubosiella* (*p* = 0.0002; **Additional file 3b**) in the stool of occultly *K. pneumoniae* colonized mice. Indeed, *Erysipelotrichaceae* consistently decreased in the vast majority of occultly colonized microbiota (**Additional file 3c**), and this pattern could be attributed to decreases in *Dubosiella* (**Additional file 3d**).

To determine the reproducibility of these changes in the microbiome, we repeated the occult colonization experiment, and this time, performed shotgun metagenomic sequencing to obtain species-level resolution. Taxonomic classification was performed using Kraken2 [35] and the Genome Taxonomy database (GTDB) [36] as reference since it was found to be superior to RefSeq, the standard database (**Additional file 4a**). First, we looked at beta diversity and found that baseline microbiota formed a separate cluster from occultly colonized microbiota, suggesting differences in microbiome structure (**Fig. 3a**). Similar to 16S rRNA results, there was a significant reduction in the abundance of *Erysipelotrichaceae* (*p* = 0.0006; **Fig. 3b and Additional file 7**), its component species *Faecalibaculum rodentium* (*p* = 0.0003), and a trend in *Dubosiella sp004793885* (*p* = 0.071) and *Dubosiella newyorkensis* (*p* = 0.071) (**Fig. 3c and Additional file 8**). However, while the relative abundance of *Erysipelotrichaceae* dropped in nearly every occultly colonized animal (**Fig. 3d**), control mice that were mock gavaged with PBS showed consistently low levels of *Erysipelotrichaceae* at both time points. The difference in starting levels could be partially explained by the larger sample size for the experimental animals (N=28) compared to control animals (N=8) but likely also reflects natural variation in the abundance of this taxon in the mouse microbiome.

**Fig. 3.**
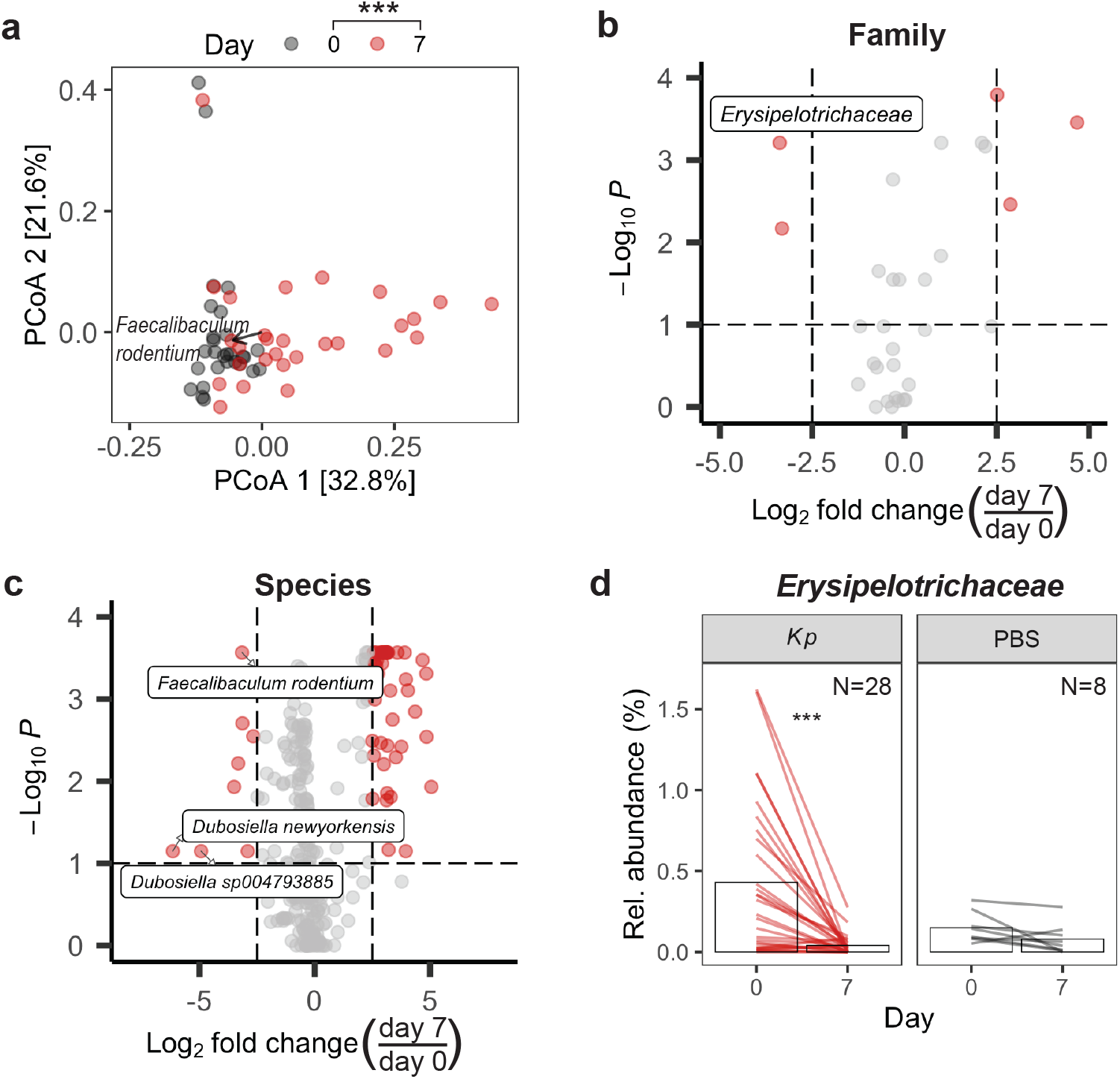
The gut microbiome is perturbed following occult colonization with *K. pneumoniae*. **a** Bray-Curtis principal coordinates analysis (PCoA) of microbiota at day 0 and day 7 post-gavage. N=28. **b, c** Volcano plots showing adjusted *p*-value versus fold change of relative abundances of taxa at the (**b**) family and (**c**) species level between the 2 timepoints. **d** Relative abundance of *Erysipelotrichaceae.* Bars indicate mean abundance. Mock gavage with PBS served as the control. N=8. *p*-values were calculated using (**a**) PERMANOVA or (**b**-**d**) paired Wilcoxon signed-rank tests with Benjamini-Hochberg correction. \*\*\**p*-value < 0.001.

### Vancomycin-induced outgrowth is incomplete and predicted by the functional potential of the microbiome

We observed in the single drug experiments (**Fig. 2b**) that vancomycin treatment resulted in *K. pneumoniae* outgrowth in only 40% of animals, whereas the 4-drug cocktail or ampicillin resulted in 100% of animals showing outgrowth. This observation led us to hypothesize that the microbiota at day 7, just before vancomycin treatment, were taxonomically or functionally different between susceptible and non-susceptible mice. First, we reproduced the stochastic outgrowth phenotype with a larger sample size (**Fig. 4a**). Next, we investigated taxonomic composition using shotgun metagenomic sequencing. At the genus level, we found a statistically significant enrichment of *Akkermansia* in non-susceptible mice at day 7 (*p* = 0.043) (**Additional file 4b**); however, significance was not observed for any specific species (lowest *p* = 0.156) (**Additional file 4c**). As this was inconclusive, we performed a second metagenomic analysis to identify enriched gene functions (**Additional file 4d**). We constructed a machine learning classifier to measure the predictive value of functional features for susceptibility. This model achieved an area-under-the-curve (AUC) score of 0.939 (highest score = 1) for distinguishing susceptible microbiota from nonsusceptible microbiota (**Fig. 4b**). The model chose 4 genes as the optimal marker set (**Fig. 4c**). The gene with the largest effect size encoded polyribonucleotide nucleotidyltransferase, which is involved in mRNA degradation. Another gene in the marker set encoded 50S ribosomal protein L16, which is involved in mRNA translation. Both genes were enriched in nonsusceptible microbiota (**Fig. 4c**). Thus, it appears, from a gene-based perspective, that functions related to mRNA, particularly its degradation, are predictors of nonsusceptibility.

**Fig. 4.**
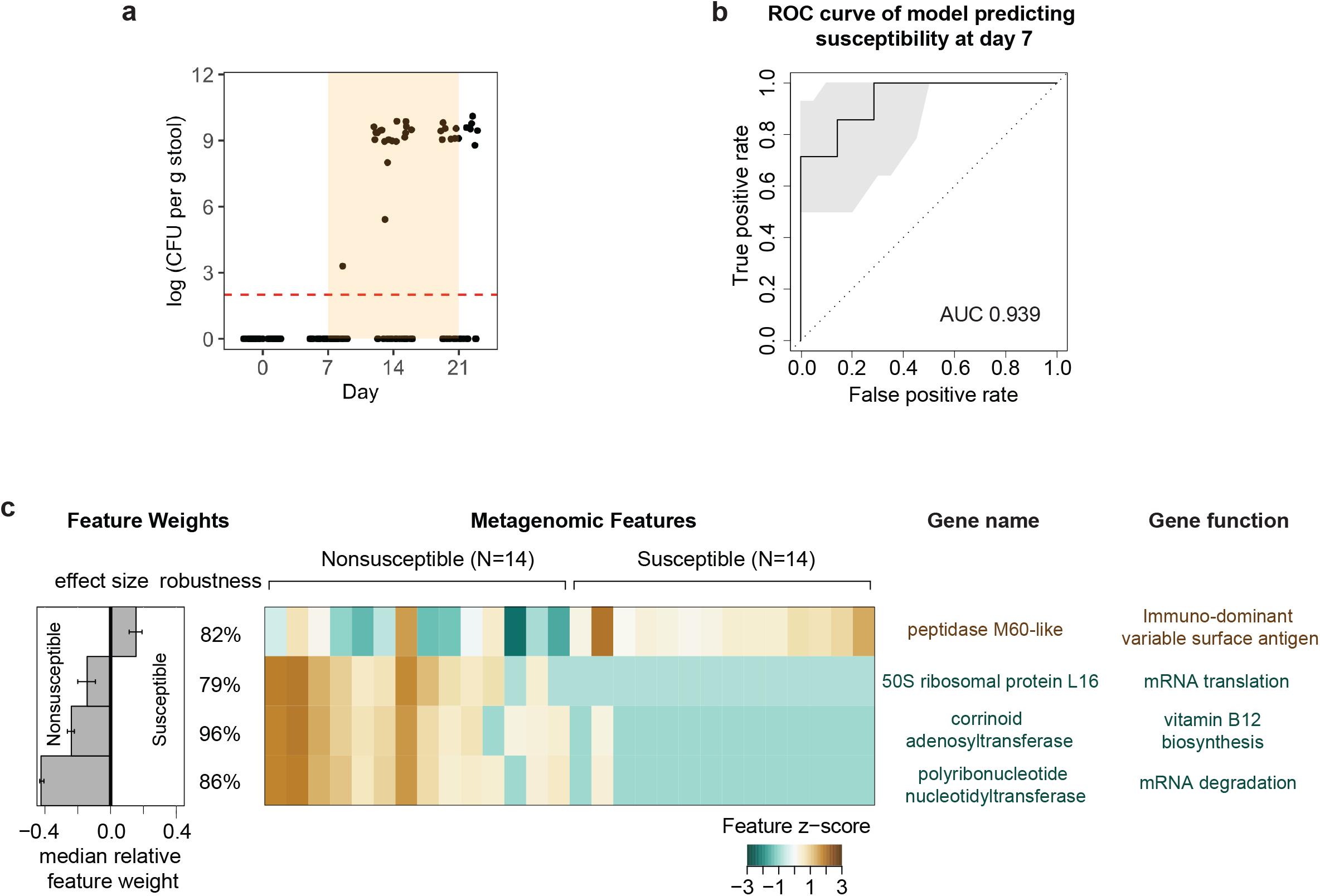
Microbiota gene functions related to mRNA degradation predict nonsusceptibility to *K. pneumoniae* outgrowth. **a** *K. pneumoniae* CFU for vancomycin-induced outgrowth in a repeat experiment using the design shown in Fig. 2a. Outgrowth and no-outgrowth were evenly divided post-antibiotic (N=14 each). **b** ROC curve for lasso machine learning model predicting susceptibility status at day 7. ROC, receiver operating characteristic; AUC, area under the curve. **c** Interpretation of the lasso machine learning model showing feature effect sizes and robustness, heatmap of top selected features, and gene names and functions.

### Vancomycin treatment selects for mutations in a *K. pneumoniae* transcriptional regulator of xylose metabolism

Up to this point, we had focused on characteristics of the microbiome that contribute to occult colonization and antibiotic-induced outgrowth. We next sought to perform a complementary experiment looking for genes in *K. pneumoniae* that contribute to colonization and growth. To this end, we passaged *K. pneumoniae* under antibiotic selection in mice to identify genes important for its outgrowth. Our rationale was that acquired mutations would elucidate the landscape of selective pressures in the antibiotic-disrupted gut, allowing us to predict what metabolites are important for outgrowth and susceptibility. Specifically, we evolved *K. pneumoniae* by iterative passages through the gut of vancomycin-treated mice (**Fig. 5a**). At the 8^th^ iteration, 3 evolved *K. pneumoniae* strains from each lineage were picked for whole-genome sequencing and analyzed for SNP variants (**Additional file 5a**). Notably, 5 out of the 6 lineages contained an isolate with a missense variant in the xylose-binding domain of the *xylR* gene (**Fig. 5b**). XylR is a transcriptional activator of the xylose operon. Structurally, XylR contains a xylose-binding domain that is critical to its transcriptional activity [37], and we noted that all five *xylR* SNP mutations were located in this exact domain, with one SNP being precisely located in a residue (Arg240) critical for functional activity [37]. Xylose can be fermented to provide energy for growth [38]. Thus, it is possible that reduced xylose may be a selective pressure in the antibiotic-treated gut causing *K. pneumoniae* non-susceptibility.

**Fig. 5.**
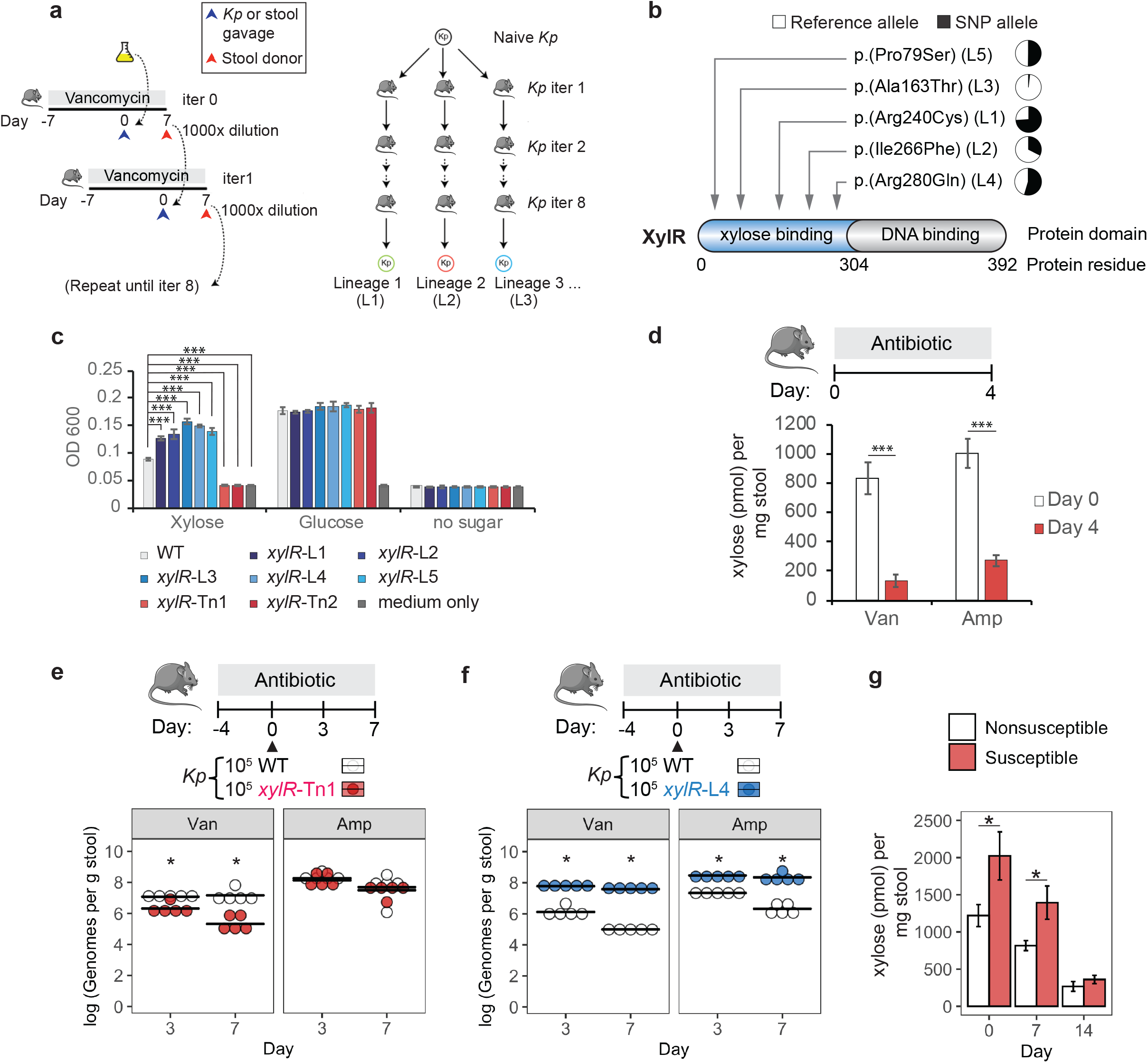
Pre-antibiotic xylose level is associated with susceptibility to *K. pneumoniae* outgrowth. **a** Schematic showing passage of *K. pneumoniae* under antibiotic selection. Mice were pre-treated with vancomycin for 7 days, orally gavaged with 10^5^ CFU of *K. pneumoniae* and maintained on vancomycin for 7 more days to allow *K. pneumoniae* growth. Stool was diluted 1000-fold to achieve ~10^5^ *K. pneumoniae* CFU, and gavaged into another vancomycin pre-treated mice. Such transfer of stool microbiota was performed for 8 iterations into singly housed mice to avoid any cross-contamination. *K. pneumoniae* colonies were cultured from the stool of the 8^th^ iteration mouse. Colonies from independent lineages were whole-genome sequenced. Plate swipes containing aggregate pools of *K. pneumoniae* cultured directly from stool were also sequenced for each lineage. **b** Summary of XylR mutations derived from 5 separate lineages together with their allelic frequencies obtained from plate swipes. **c***In vitro* anaerobic growth assays of evolved strains *xylR*-L and transposon strains *xylR*-Tn in minimal media containing xylose or glucose after 24 hours. N=4. **d** Levels of xylose in stool before and after vancomycin or ampicillin treatments. N=10. **e**, **f***In vivo* competition between (**e**) the wild type and *xylR*-Tn1 transposon strain, or (**f**) the wild type and *xylR*-L4 evolved strain in mice undergoing vancomycin or ampicillin treatment. Mice were orally gavaged with 10^5^ CFU of each *K. pneumoniae* strain at day 0 (triangle). Strain levels at days 3 and 7 were quantified by qPCR using specific primers to each *xylR* allele. N=5. **g** Levels of xylose in stool at day 0, 7 or 14. Stool was obtained from the mice shown in Fig. 4a. N=13-14. For (**c**), one-way ANOVA was performed followed by post-hoc Dunnett’s test comparing each evolved or Tn strain to the wild type on each sugar. For (**d**, **g**), a two-tailed t-test was used. For (**c**, **d** and **g**), error bars show SEM. For (**e**, **f**), a Mann-Whitney U-test was used to compare evolved or mutant strains to the wild type at each time-point. \**p*-value < 0.05, \*\*\**p*-value < 0.001.

We next tested the 5 independently evolved *xylR* mutants for altered antibiotic resistance but found no difference in the minimum inhibitory concentrations (MICs) of vancomycin and ampicillin compared to the wild type (**Additional file 1a**). This suggests that the *xylR* variants did not confer altered resistance to vancomycin or ampicillin. We next assessed whether the variants alter xylose utilization by comparing the growth of the wild type and 5 *xylR* mutants in minimal media with xylose or glucose as the sole source of carbon (**Fig. 5c**). While the wild type and all 5 mutants showed similar growth on glucose, the mutants showed significantly higher growth on xylose (*p* < 0.001), suggesting that the SNPs are gain-of-function mutations that enhance the utilization of xylose. Supporting this, loss-of-function *xylR* transposon mutants [24] were incapable of utilizing xylose as a sole carbon source (**Fig. 5c**). These data suggest that the murine evolved strains are gain-of-function with superior growth on xylose compared with the wild type.

### Xylose metabolism contributes to *K. pneumoniae* growth in the context of antibiotic therapy

Based on these data, we hypothesized that antibiotic treatment reduces the level of xylose, driving *K. pneumoniae* to evolve mutations that enhance xylose utilization. To explore this idea, we quantified the amount of xylose in stool before and after antibiotic treatment, and indeed found significantly reduced levels of xylose in both vancomycin-treated (*p* = 6 x 10^−5^) and ampicillin-treated stool (*p* = 2 x 10^−5^) (**Fig. 5d**).

The adaptation of *K. pneumoniae* to a changing xylose supply suggests the importance of xylose metabolism for *K. pneumoniae* growth. We tested this by gavaging mice with equal amounts of wild-type *K. pneumoniae* and a *xylR* transposon knockout. We found lower levels of the *xylR* transposon strain compared to the wild-type strain in vancomycin-treated mice (*p* = 0.012 at day 7), suggesting that the induction of xylose utilization via XylR provides a competitive growth benefit (**Fig. 5e**). To test whether the *xylR* variants in the evolved strains provide enhanced growth *in vivo*, we colonized vancomycin-treated mice with equal amounts of the wild-type strain and an evolved strain (**Fig. 5f**), the latter of which possessed a SNP only in the *xylR* gene (**Additional file 5a**). We found that the evolved strain significantly outcompeted the wild-type strain (*p* = 0.012). To determine if the growth superiority of the evolved strain is limited to vancomycin-treated mice, we performed a similar experiment in ampicillin-treated mice (**Fig. 5f**) and observed that the evolved strain also showed enhanced growth compared to wild-type strain (*p* = 0.012). These results demonstrate that xylose metabolism is important for the growth of *K. pneumoniae* in the context of vancomycin and ampicillin receipt.

### Elevated intestinal xylose is a marker of susceptibility to *K. pneumoniae* outgrowth

These results prompted us to revisit the incomplete vancomycin-induced outgrowth of occultly colonized mice (**Fig. 4a**). We hypothesized that xylose may be associated with susceptibility prior to antibiotic treatment. We found that xylose was significantly higher in susceptible compared to nonsusceptible stool at day 7 (*p* = 0.026) and, interestingly, at the day 0 baseline as well (*p* =0.038; **Fig. 5g**). These results support the use of pre-antibiotic xylose levels as a marker of susceptibility.

## Discussion

In this work, we present an optimized mouse model for use in occult gut colonization studies. It is clinically relevant, uses common immune-competent mice without antibiotic pre-treatment, and is stable and reproducible. We observed consistent antibiotic-induced outgrowth patterns over a total of N=330 mice, including incomplete vancomycin-associated outgrowth (**Additional file 5b**). We successfully reproduced these essential features in a different mouse facility (data not shown). Similar to previous mouse models of *K. pneumoniae* colonization [39], we chose as our occult colonizer a genetically tractable, human-associated ST258 strain of *K. pneumoniae*, but demonstrated that other *Enterobacteriaceae* may also be used. Using this system, we observed that occult colonization remodels the microbiota and that genes regulating mRNA stability and xylose metabolism influence subsequent antibiotic-triggered outgrowth. Previous studies on occult colonization have shown the ability of exogenous strains to adhere to the intestinal mucosa [40], but mechanistic understanding is fairly limited. In other studies involving a somewhat similar process of persistent colonization, it was found that some of the important factors are iron scavenging [41], utilization of sialic acid from mucus layer [42], and bacteriocins to compete against other strains [43]. It may be interesting to see whether these factors are also important in occult colonization. Similar approaches could be conducted for other strains of interest to study their occult properties.

We studied the impact of low-level *K. pneumoniae* colonization on the microbiome and observed a reduced abundance of *Erysipelotrichaceae*. Due to the large variability in baseline taxa abundances, the experiments presented here did not have the power to identify marker taxa of occult colonization from the complex native gut microbiome. Future experiments using more timepoints, additional animals or a defined gut community could be used to identify these marker taxa.

We also investigated the functional potential of susceptible microbiota and found genes regulating mRNA stability as a predictor of susceptibility. Response to stress caused by antibiotics involves changes in the transcriptional state of the bacteria [44], particularly elevated synthesis of selected stress response proteins [45, 46]. To achieve this regulation, it is thought that keeping a low general mRNA abundance allows crucial mRNA encoding stress-related proteins to gain access to ribosomes for translation [47, 48]. Our metagenomic analysis of nonsusceptible microbiota potentially supports this paradigm of translational resource reallocation, in which mRNAs not related to stress responses are selectively degraded. Our findings of enhanced mRNA degradation, along with enhanced ribosomal capacity, may allow the microbiota to better cope with antibiotic stress and resist *K. pneumoniae* outgrowth. It is a goal of future studies to probe how the dynamics and regulation of mRNA stability affect susceptibility.

We also explored the impact of small molecules on *K. pneumoniae* outgrowth and identified xylose as a potential marker molecule. An advantage of using xylose as a marker of susceptibility is that xylose is also present in human stool as a breakdown product of dietary fiber [49, 50]. Plant-derived dietary xylose is a major carbon source for the gut microbiota, with different microbiota members playing roles in branched-chain xylan degradation and the cross-feeding of other members of the community [51–53]. We found that *K. pneumoniae* relies on this source of free xylose for growth, in agreement with another recent study [54], and we went a step further to link xylose levels with susceptibility. Another strength of xylose as a marker of susceptibility is that the level is relatively stable over time, as both day 0 and day 7 in our model showed significant differences in xylose abundance between susceptible and nonsusceptible mice. Moreover, xylose abundance can be measured using a commercial kit in a matter of hours and therefore does not require costly metabolomics. Future work to improve health outcomes may investigate how to therapeutically reduce levels of free xylose, such as by catalyzing its fermentation to short chain fatty acids [38], which have numerous health benefits for the host [55, 56].

A limitation of the work is that xylose abundance is not as effective for predicting susceptibility to ampicillin-induced outgrowth. This suggests that during ampicillin treatment, xylose may be compensated for by other carbon sources, such as fucose or sialic acid [42, 57], thus allowing outgrowth regardless of xylose availability. Moreover, the microbiome of mouse and human have considerable differences [58], hence further work is needed to confirm whether our findings on mouse microbiome extend to human microbiome.

## Conclusion

CRE infections are responsible for high patient mortality and comorbidities and are difficult to treat because of their resistance to broad classes of antibiotics. The epidemiology of CRE transmission is complicated to track because of frequent patient transfers between healthcare facilities and increasing global travel including leisure or medical tourism. Occult CRE colonization compounds this difficulty by allowing an undetectable reservoir in carriers and causing an underestimation of CRE prevalence in healthcare systems and in the community. It is clinically important to understand which individuals are predisposed to outgrowth in order to estimate the risks of different antibiotic treatments. We identified two susceptibility factors, mRNA stability pathways and intestinal xylose abundance, using two complementary approaches. These results provide taxonomic, functional and genetic insight into CRE occult colonization and outgrowth, and pave the way for mechanistic studies to come.

## Methods

### Mouse husbandry

C57BL/6N wild-type mice were bred under specific-pathogen-free conditions at an American Association for the Accreditation of Laboratory Animal Care-accredited animal facility at the NHGRI and housed in accordance with procedures outlined in the Guide for the Care and Use of Laboratory Animals. All experiments were performed under an animal study proposal approved by the NHGRI Animal Care and Use Committee. Cages, water and feed were pre-sterilized by autoclaving before use. Gender- and age-matched mice between 7 to 12 weeks old were used in all experiments. Cross-contamination between cages was minimized during cage handling by frequent changes of gloves and disinfection of hands with 200 ppm chloride dioxide. For all experiments, mice were singly housed to prevent transmission of *K. pneumoniae* by coprophagy.

### Antibiotics

Mice were treated with antibiotics in drinking water for 2 weeks. Antibiotics used were 0.5 g/L ampicillin (Sigma), 0.5 g/L vancomycin (Letco), 0.5 g/L neomycin (Sigma), and 0.5 g/L metronidazole (Sigma), 0.2 g/L ciprofloxacin (Sigma), 0.1 g/L azithromycin (Sigma), 0.1g/L rifaximin (Sigma). The antibiotic cocktail was 0.5 g/L ampicillin, 0.5 g/L vancomycin, 0.5 g/L neomycin, and 0.5 g/L metronidazole.

### Inoculation of *K. pneumoniae*, *E. coli* and *E. cloacae*

*K. pneumoniae*, *E. coli* and *E. cloacae* were cultured overnight in LB at 37°C under aerobic conditions with shaking at 200 rpm. Unless otherwise stated, mice were orally gavaged with 10^5^ CFU, and all experiments involving *K. pneumoniae* used MKP103 strain as the wild type. MKP103 is a derivative of ST258 KPNIH1 in which the *bla*_KPC_ gene was deleted [24]. All other *K. pneumoniae*, *E. coli* and *E. cloacae* strains used in the study harbor the *bla*_KPC_ gene, are resistant to carbapenem and ampicillin, and are derived from human sources. More details of each strain are given in Additional file 6b.

For *in vivo* competition of *K. pneumoniae* strains, mice received vancomycin or ampicillin for 3 days prior to being orally gavaged with 10^5^ CFU of an evolved/ transposon strain and 10^5^ CFU of MKP103. Mice were maintained on antibiotics for 7 more days.

### Collection of stool and cecal content

Stool pellets were collected from mice at days 0, 7, 14, and 21 unless otherwise stated. For each time point, a fresh pellet was used for CFU counting, while another pellet was snap-frozen on dry ice and stored at −80°C for 16S rRNA or metagenomic sequencing. Cecal contents were obtained by making a cut along the cecal pouch. Small intestinal contents were flushed out with PBS using a syringe and oral gavage tube. Colonic contents were gently massaged out from either end of the colon using a pair of forceps. Mesenteric lymph nodes and livers were homogenized through 70 μm strainers prior to plating. Whole blood was collected into blood plasma tubes and plated directly.

### Quantification of CFU

To determine CFU for *K. pneumoniae* MKP103, samples were serially diluted and plated onto LB agar containing 100 μg/ml vancomycin, 100 μg/ml metronidazole, 100 μg/ml neomycin and 50 μg/ml ampicillin. Plates were incubated for 18 hours at 37°C under aerobic conditions. Negative control plating of wild-type stool typically showed zero colonies. 5 to 10 randomly selected colonies were verified per experiment with PCR using *K. pneumoniae*-specific primers (see Additional file 6a for primer sequences).

### *In vitro* growth assays

*In vitro* growth assays were performed in M9 minimal medium (BD) supplemented with 20 mM xylose or glucose (Sigma). Wild-type and *xylR* evolved/ transposon strains were first cultured at 37°C for 24 hours in an anaerobic box (AnaeroPack system, Mitsubishi Gas Chemical Company) with disposable single-use anaerobic sachets (GasPak EZ Anaerobe Container System, BD). Then, strains were washed and resuspended at 10^5^ CFU/ml in M9 with 20 mM sugar and incubated anaerobically in 96-well plates in 100 μl final volumes for 24 hours. After 24 hours, the OD at 600 nm was measured with a plate reader (Biotek).

### Xylose quantification

Free xylose levels were quantified using a colorimetric-based D-xylose assay kit (Megazyme). Stool samples were weighed, resuspended in 1 ml ddH2O, vortexed for 30 seconds, and then centrifuged at 13,000 g for 2 min to remove large particles. Next, supernatants were transferred to new tubes, and the manufacturer’s instructions were closely followed to measure xylose levels, including the step of Carrez purification to remove protein inhibitors.

### Determinaton of minimum inhibitory concentrations

MICs for vancomycin, metronidazole, ampicillin, ciprofloxacin and azithromycin were determined using MIC test strips (Liofilchem) according to the manufacturer’s instructions. MICs for rifaximin and neomycin were determined for two-fold serial dilutions of the antibiotic from 0.5 μg/ml to 256 μg/ml. Bacteria were inoculated at 5 x 10^4^ CFU in 100 μl in 96-well plates. After overnight incubation at 37°C, OD600 was measured. MIC was defined as the lowest concentration with no bacterial growth.

### DNA extraction

DNA was extracted from stool using the DNeasy Powersoil Kit (Qiagen) according to the manufacturer’s instructions except that, prior to any processing, 200 μl of the solution in the Powerbead tube was replaced with 200 μl of phenol:chloroform:isoamyl alcohol (25:24:1, Thermo Fisher).

### Quantitative PCR

qPCR was used to quantify levels of carbapenemase producing clinical strains and *xylR* mutants. 2 μl of the DNA that was extracted from stool was used in 10 μl reactions with Powerup SYBR Green PCR master mix (Applied Biosystems) in 384-well plates. qPCR reactions were run using a Quant Studio 6 Flex (Applied Biosystems). Primer sequences are provided in Additional file 6a.

### 16S rRNA sequencing

To resolve *K. pneumoniae* at the genus level, the V1-V3 region of the 16S rRNA gene was sequenced instead of the V4 region. V1-V3 sequencing libraries were generated by PCR using barcoded primers flanking V1 (27F, 5’-AGAGTTTGATCCTGGCTCAG-3’) and V3 (534R, 5’-ATTACCGCGGCTGCTGG-3’). PCR amplicons were purified using Agencourt AMPure XP beads (Beckman Coulter), quantified using the Quant-iT dsDNA high-sensitivity assay kit (Thermo Fisher), pooled in equimolar amounts, and re-purified with the MinElute PCR purification kit (Qiagen). Sequencing was performed on an Illumina MiSeq at the NIH Intramural Sequencing Center (NISC).

### 16S rRNA sequencing analysis

Taxonomic classification was performed on the first 200 bp (quality score >30) of read 1 using the R package dada2 v1.12.1 [59] against the Silva database (release 132) [60]. The dada2 output containing 1,186 amplicon sequence variants (ASVs) was imported into the R package Phyloseq v1.28.0 [61] for downstream analysis. Alpha diversity (number of ASVs) was calculated using the estimate_richness function with option ‘Observed’ in Phyloseq. Volcano plots were generated using the R package Enhanced volcano [62]. Differential abundance testing between timepoints was performed on relative abundances using paired Wilcoxon signed-rank test with Benjamini-Hochberg correction.

### Shotgun metagenomics sequencing

Libraries were prepared as described [63] using the Illumina Nextera XT kit for paired-end (2 x 150bp) sequencing on an Illumina NovaSeq at NISC. Reads were processed by trimming adapters using cutadapt [64], removing low quality reads using prinseq-lite [65] with option ‘-min_qual_mean 20’ and removing host (mouse; GRCm39) reads. A total of 100 samples were sequenced, resulting in 5.98 billion non-mouse, quality-filtered paired-end reads (average 59.8 million reads per sample).

### Shotgun metagenomics community composition analysis

Metagenomic reads were taxonomically classified using Kraken2 [35] with option ‘--confidence 0.1’. Taxonomic classification was performed using either the built-in standard RefSeq database or a custom GTDB database (release 95) [36] built using the Struo workflow [66]. A total of 27,130 GTDB bacterial and archaeal genomes from a total of 194,600 were selected to build the database using the following criteria: > 50% completeness, < 5% contamination and being a representative genome. No fungi, viral or protozoa genome were included. Taxonomy was further speciated using Bracken [67] with option ‘-t 1000’ to remove low abundance species. In total, 706 species were classified. Bracken output was converted into the BIOM format using Kraken-biom (https://github.com/smdabdoub/kraken-biom) and imported into Phyloseq for downstream analyses. To generate Bray-Curtis PCoA plots, ASV abundances were variance-stabilized using Hellinger transformation via the ‘decostand’ command in the R package vegan [68] and plotted using the ‘ordinate’ command in Phyloseq. Differential abundance testing between unpaired samples was performed in MaAsLin2 [69] using the linear model method with Benjamini-Hochberg correction.

### Shotgun metagenomics functional analysis

Metagenomic datasets were assembled using SPAdes v3.14.0 [70] with the option --meta. Protein-coding sequences (CDS) for assembled contigs were predicted and annotated with Prokka v1.14.0 [71] with the option --metagenome. Protein clustering was performed using the ‘linclust’ function in MMseqs2 [72] with options ‘--cov-mode 1 -c 0.8’ (minimum coverage threshold of 80% the length of the shortest sequence) and ‘--kmer-per-seq 80’. The ‘--min-seq-id’ option was set at 0.95 to generate the catalogues at 95% protein identity. A total of 467,873 proteins were obtained in the catalogue. Functional characterization of all protein sequences was performed with eggNOG-mapper v2 [73]. The clustered protein sequences were converted to a DIAMOND database [74]. The metagenomic reads were aligned against the protein catalogue using DIAMOND blastx with the options --id 90, --evalue 1e-6, -k 1, --max-hsps 1, and --unal 0. Proteins were filtered to remove low-abundance counts and converted to relative abundance by normalizing to total reads. Machine learning of proteins was performed using SIAMCAT v1.10.0 [75] with leave-one-out cross-validation (i.e., 28-fold cross-validation for n=28 samples) and the ‘lasso’ method. Robustness refers to the proportion of machine learning models that have incorporated the feature weights. Heatmap was plotted after converting relative abundance to z-score.

### Whole-genome and metagenomic sequencing of evolved strains

The evolution strategy described in Figure 5a produced 3 evolved *K. pneumoniae* isolates from each of 6 independent lineages. DNA was extracted from fresh cultures of these 18 strains on an automated platform (Promega Maxwell) as previously described [23]. To examine *K. pneumoniae* evolution on a metagenomic level, DNA was also extracted from pooled *K. pneumoniae* colonies cultured directly from the stool of each mouse lineage. Culture plates with 100-150 colonies were swiped together (plate swipes), and stored in 15% glycerol at −80°C until DNA extraction. Illumina libraries were created using Nextera library chemistry. Sequencing was performed on an Illumina MiSeq at the NIH Intramural Sequencing Center.

### SNP calling

SNP calling for individual *K. pneumoniae* isolates was performed by aligning the whole-genome sequencing reads for these isolates to the KPNIH1 reference genome and calling SNPs using SNIPPY v3.2 with default settings (https://github.com/tseemann/snippy). Identified SNPs were validated by Sanger sequencing. In addition to the SNPs reported in Additional file 5a, there were 3 SNPs inherent to MKP103 that presumably occurred during the growth and selection for deletion of the *bla*_KPC_ gene from KPNIH1. These 3 SNPs, which were present in all evolved strains, included a missense SNP in KPNIH1_08215 (hypothetical protein), a synonymous SNP in KPNIH1_09140 (lipid transporter ATP-binding/permease), and a synonymous SNP in KPNIH1_09495 (gamma-glutamylputrescine synthetase). SNPs were named according to the Human Genome Variation Society (HGVS) nomenclature [76].

For metagenomic SNP calling, allelic frequencies of SNPs from the sequencing of plate swipes were determined using bam-readcount (https://github.com/genome/bam-readcount). Briefly, reads were aligned to the KPNIH1 reference genome using bowtie2 [77], and the resulting sam file was converted into a bam file, sorted, indexed, and input into bam-readcount. Additionally, the known SNP locations for each lineage that had been observed using SNIPPY for isolate genomes were also input into bam-readcount to extract allelic frequencies for these SNPs in each plate swipe. Sequencing depth of the 5 *xylR* SNP locations was between 209 and 293 reads.

### Statistics

For normally distributed data, a two-tailed t-test was used to compare 2 groups, whereas multiple groups were compared using one-way ANOVA followed by a post-hoc Dunnett’s test to compare treatment groups against a control group. For non-normally distributed data, an unpaired Mann-Whitney U test or paired Wilcoxon signed-rank test was used to compare 2 groups, whereas multiple groups were compared using the Kruskal-Wallis H test followed by pairwise Mann-Whitney U tests to compare each treatment group against a control group. Correction for multiple testing was performed using the Benjamini-Hochberg method when the number of tests exceeded 20; otherwise, Bonferroni correction was used. Significance for categorical variables (e.g. outgrowth versus no outgrowth) was determined using Fisher’s exact test with Hommel correction for multiple testing against a control group. PERMANOVA was calculated using the Adonis function in the R package vegan. Normality was assessed using the Shapiro-Wilk test. All statistical analyses were performed in R.

---

**Additional file 1 a** Table showing minimum inhibitory concentration (MIC) of antibiotics on wild-type KPNIH1 derivative (MKP103), evolved strain (*xylR*-L4) and transposon strain (*xylR*-Tn1). ND, not determined. **b** Table showing antibiotics used in study, their drug class, mechanisms of actions and target bacteria.

**Additional file 2** KPC-producing strains of *K. pneumoniae, E. cloacae* and *E.coli* occultly colonize the gut until robust outgrowth by ampicillin. Mice were orally gavaged with 10^5^ CFU of each strain or PBS at day 0. Ampicillin was given from day 7 to day 21. Levels of *Enterobacteriaceae* in stool at the stated time-points.

Measurement was done by qPCR using primers at *bla*_KPC_ gene and Ct values were converted to number of genomes using a standard curve made from dilutions of known DNA mass. The red dotted line marks the limit of detection. N=4-9. Statistics was performed using Fisher’s exact test with Hommel correction comparing each strain to PBS control on each time-point. \**p*-value < 0.05, \*\**p*-value < 0.01, \*\*\**p*-value < 0.001

**Additional file 3 a**, **b** Volcano plots showing adjusted *p*-value versus fold change of relative abundances of taxa at (**a**) family level and (**b**) genus level between day 0 and day 7. **c**, **d** Relative abundances of (**c**) *Erysipelotrichaceae* and (**d**) component genus *Dubosiella*. Bars show mean abundance. N=64. For (**a**-**d**), *p*-value is calculated by paired Wilcoxon signed-rank test with Benjamini-Hochberg correction. \*\*\**p*-value < 0.001.

**Additional file 4 a** Percentage of reads with a Kraken2 taxonomic classification using standard RefSeq or GTDB databases. Error bars denote SEM. **b** Relative abundance of the only significant genus *Akkermansia* between susceptible and nonsusceptible. **c** Top 5 species that show a trend between susceptible and nonsusceptible. **d** Percentage of reads with an eggNOG functional annotation. For (**a**), *p*-value is calculated by a two-tailed t-test. For (**b**) and (**c**), *p*-value is calculated using the linear model in MaAsLin2 and adjusted by Benjamini-Hochberg. \**p*-value < 0.05, \*\*\**p*-value < 0.001.

**Additional file 5 a** Table showing mutations in evolved strains. A representative strain is displayed per lineage. **b** Overall outgrowth statistics of the stated antibiotics from all the mice used in the project.

**Additional file 6 a** Primer sequences. **b** Bacterial strains

**Additional file 7** Abundance, log2 fold change, and *p*-values of taxa between day 0 and day 7 at family level. Supporting information for fig. 3.

**Additional file 8** Abundance, log2 fold change, and *p*-values of taxa between day 0 and day 7 at species level. Supporting information for fig. 3.

## Abbreviations

CRE: carbapenem-resistant *Enterobacteriaceae*
*Kp*: *Klebsiella pneumoniae*
KPC: *K. pneumoniae* carbapenemase
MIC: minimum inhibitory concentration
PCoA: principal coordinate analysis
ROC: receiver operating characteristic
AUC: area under the curve
CFU: colony-forming unit
SI: small intestine
MLN: mesenteric lymph node
GTDB: Genome Taxonomy database
ASV: amplicon sequence variant
SNP: single nucleotide polymorphism

## Declarations

### Ethics approval and consent to participate

The animal studies were approved by the NHGRI Animal Care and Use Committee.

### Consent for publication

Not applicable.

### Availability of data and material

The datasets analyzed in this paper are deposited at BioProject: PRJNA687411.

### Competing interests

The authors declare that they have no competing interests.

### Funding

This work was supported by the Intramural Research Programs of NHGRI and NIAID.

### Authors’ contributions

CKS and JAS designed the study. CKS, JAS, SC, and AS wrote the manuscript. CKS performed experiments. CKS and SSK analyzed shotgun metagenomic data. CKS and DMP analyzed 16S rRNA data. CKS and SC analyzed whole-genome sequencing data. AS, CD assisted with microbiology experiments. MC assisted with evolution experiments. NB and YB provided immunology support. AA and RDF assisted with functional metagenomic analyses. All authors reviewed and edited the manuscript.

## Acknowledgements

We thank Qiong Chen for making metagenomics sequencing libraries, Shih-Queen Lee-Lin and Clay Deming for making 16S rRNA sequencing libraries, Ryan C. Johnson for advice on SNP analysis, Jessica LeGrand for technical support with mouse work, Ivan Vujkovic-Cvijin for intellectual discussions, Paul Juneau for consultation on statistics, Tara N. Palmore for suggestions on clinical relevance, Paul E. Carlson lab for microbiology support, the Segre lab and Heidi Kong lab for general discussions, the NHGRI and NIAID animal facilities for mouse husbandry, and the NIH High-Performance Computation Biowulf Cluster (http://hpc.nih.gov) for the computational resources.

## Notes

### Competing Interest Statement

The authors have declared no competing interest.

