## Additional File 1 for "A mouse model of occult intestinal colonization demonstrating antibiotic-induced outgrowth of carbapenem-resistant *Enterobacteriaceae*"

**a**

| | Antibiotic | MKP103 | xylR-L4 | xylR-Tn1 |
|---|---|---|---|---|
| *1* | Ampicillin | >256 | >256 | >256 |
| *2* | Vancomycin | >256 | >256 | >256 |
| *3* | Neomycin | 256 | ND | ND |
| *4* | Metronidazole | >256 | ND | ND |
| *5* | Ciprofloxacin | >32 | ND | ND |
| *6* | Azithromycin | >256 | ND | ND |
| *7* | Rifaximin | >256 | ND | ND |

ND = not determined

**b**

| | Antibiotic | Drug class | Mechanism | Target |
|---|---|---|---|---|
| *1* | Cocktail (AMNV) | See Below | See Below | See Below |
| *2* | Ampicillin (A) | beta-lactam, penicillin | Inhibits cell wall synthesis | Many gram-positive and some gram-negative bacteria |
| *3* | Metronidazole (M) | Nitroimidazole | Destabilizes DNA | Anaerobes |
| *4* | Neomycin (N) | Aminoglycoside | Inhibits protein synthesis | Gram-positive and -negative bacteria |
| *5* | Vancomycin (V) | Glycopeptide | Inhibits cell wall synthesis | Gram-positive bacteria |
| *6* | Ciprofloxacin | Fluoroquinolone | Inhibits DNA gyrase | Some gram-positive and many gram-negative bacteria |
| *7* | Azithromycin | Macrolide | Inhibits protein synthesis | Many gram-positive and some gram-negative bacteria |
| *8* | Rifaximin | Rifamycins | Inhibits RNA polymerase | Gram-positive and -negative bacteria |
