## Supplementary figures and images for "A mouse model of occult intestinal colonization demonstrating antibiotic-induced outgrowth of carbapenem-resistant *Enterobacteriaceae*"

### Additional File 2

Additional file 2

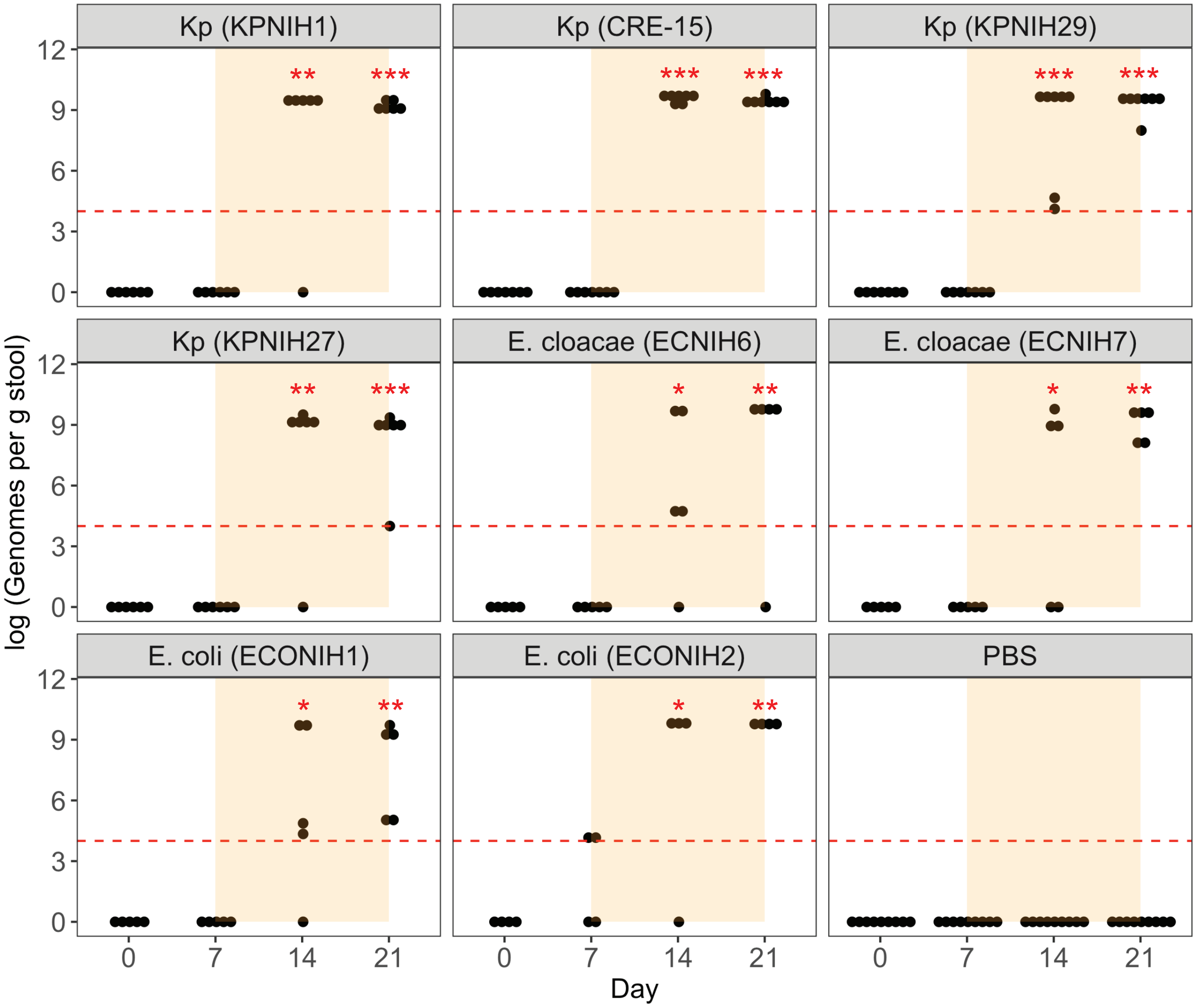

### Additional File 3

Additional file 3

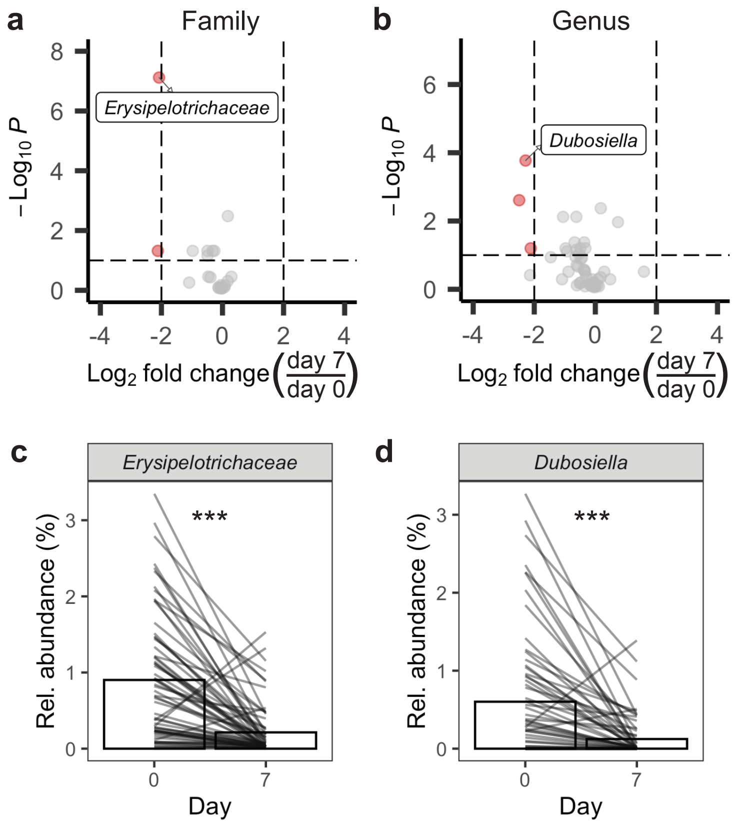
