## Additional File 4 for "A mouse model of occult intestinal colonization demonstrating antibiotic-induced outgrowth of carbapenem-resistant *Enterobacteriaceae*"

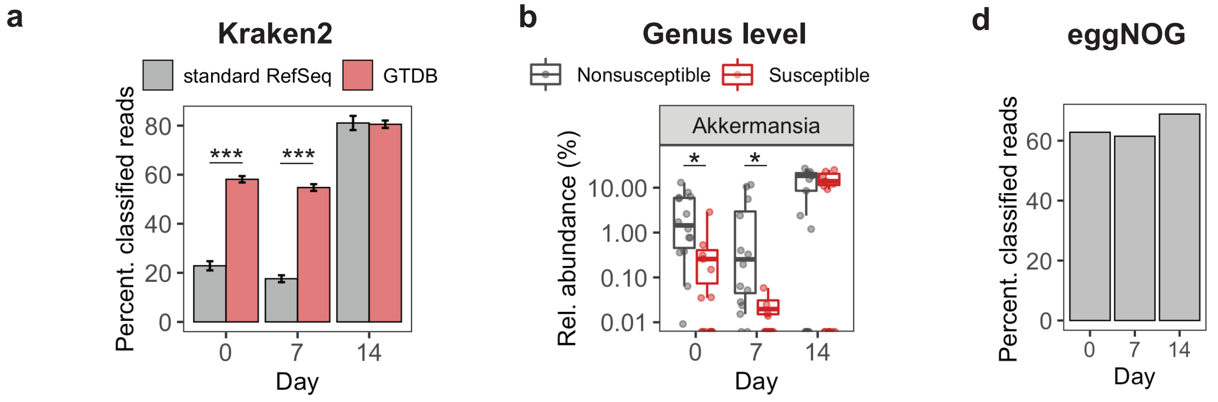


c

| | day 0 | | day 7 | |
|---|---|---|---|---|
| **Species level** | **adjusted pval** | **enriched** | **adjusted pval** | **enriched** |
| Akkermansia muciniphila A | 0.122 | nonsuscep | 0.156 | nonsuscep |
| Bacteroides ovatus | 0.122 | nonsuscep | 0.156 | nonsuscep |
| Duncaniella sp001701225 | 0.122 | suscep | > 0.2 | suscep |
| Alistipes sp003979135 | 0.122 | suscep | > 0.2 | suscep |
| Akkermansia muciniphila | 0.122 | nonsuscep | > 0.2 | nonsuscep |
