## Additional File 5 for "A mouse model of occult intestinal colonization demonstrating antibiotic-induced outgrowth of carbapenem-resistant *Enterobacteriaceae*"

**a**

| Evolved lineage | Mutation (SNP/ deletion) | Gene | Gene product |
|---|---|---|---|
| L1 | missense_variant c.163C>T p.His55Tyr | KPNIH1_02290 | 30S ribosomal protein S6 |
| | conservative_inframe_deletion c.399_404delCGGCGA p.Gly134_Asp135del | KPNIH1_18070 | porin OmpC |
| | missense_variant c.765G>T p.Met255Ile | KPNIH1_22140 | S-adenosylmethionine synthetase |
| | missense_variant c.718C>T p.Arg240Cys | KPNIH1_25345 | XylR family transcriptional regulator |
| L2 | missense_variant c.199A>G p.Thr67Ala | KPNIH1_12760 | DeoR family transcriptional regulator |
| | missense_variant c.448A>C p.Thr150Pro | KPNIH1_24015 | multidrug transporter |
| | missense_variant c.796A>T p.Ile266Phe | KPNIH1_25345 | XylR family transcriptional regulator |
| | stop_gained c.543G>A p.Trp181* | KPNIH1_26320 | phosphate ABC transporter substrate-binding protein |
| L3 | conservative_inframe_deletion c.399_404delCGGCGA p.Gly134_Asp135del | KPNIH1_18070 | porin OmpC |
| | missense_variant c.487G>A p.Ala163Thr | KPNIH1_25345 | XylR family transcriptional regulator |
| L4 | missense_variant c.839G>A p.Arg280Gln | KPNIH1_25345 | XylR family transcriptional regulator |
| L5 | missense_variant c.53T>G p.Ile18Ser | KPNIH1_03765 | two-component response regulator DpiA |
| | conservative_inframe_deletion c.399_404delCGGCGA p.Gly134_Asp135del | KPNIH1_18070 | porin OmpC |
| | missense_variant c.235C>T p.Pro79Ser | KPNIH1_25345 | XylR family transcriptional regulator |
| L6 | missense_variant c.179G>A p.Arg60His | KPNIH1_17610 | LacI family transcriptional regulator |
| | conservative_inframe_deletion c.399_404delCGGCGA p.Gly134_Asp135del | KPNIH1_18070 | porin OmpC |

**b**

| Antibiotic | No. of mice tested | Outgrowth (%) at day 21 |
|---|---|---|
| cocktail | 45 | 91.1 |
| amp | 96 | 99.0 |
| van | 141 | 50.4 |
| azm | 48 | 29.2 |
| Total | 330 | |
