## Additional File 6 for "A mouse model of occult intestinal colonization demonstrating antibiotic-induced outgrowth of carbapenem-resistant *Enterobacteriaceae*"

**a**

| Allele | Forward primer (5' to 3') | Reverse primer (5' to 3') | Notes |
|---|---|---|---|
| blaKPC | CCATCCGTTACGGCAAAAAT | TTATCACTGTATTGCACGGCG | For figure 2e and S2 |
| xylR-L4 | ACGCTCCACCGACTACCA | CCCCGACCTCCTCTTTGAAG | For figure 5f |
| xylR-WT (control for L4) | ACGCTCCACCGACTACCG | CCCCGACCTCCTCTTTGAAG | For figure 5f |
| xylR-Tn1 | GGTCCACTATATCGCCACCG | GAACACCCGAGAAAATTCATCG | For figure 5e. Use PCR elongation time = 20 sec |
| xylR-WT (control for Tn1) | GGTCCACTATATCGCCACCG | GGCATACTCGCGCTCTACC | For figure 5e. Use PCR elongation time = 20 sec |
| KPNIH1_17390 | GAATACCTTTCGCCTTGATGC | GTGGTGGGTGTGTATCGC | Specific for K. pneumoniae. |

**b**

| Bacterial strains | Description | Source |
|---|---|---|
| MKP103 | K. pneumoniae, ST258, blaKPC-, derived from KPNIH1 strain | (Ramage et al. 2017) |
| KPNIH1 | K. pneumoniae patient isolate, ST258, blaKPC+ | (Conlan et al. 2014) |
| KPNIH29 | K. pneumoniae patient isolate, ST1518, blaKPC+ | (Conlan et al. 2014) |
| KPNIH27 | K. pneumoniae patient isolate, ST34, blaKPC+ | (Conlan et al. 2014) |
| CRE-15 | K. pneumoniae patient isolate, ST45, blaKPC+ | NCBI accession: SAMN04014967 |
| ECONIH1 | E. coli patient isolate, ST648, blaKPC+ | (Conlan et al. 2014) |
| ECONIH2 | E. coli patient isolate, ST127, blaKPC+ | (Hardiman et al. 2016) |
| ECNIH6 | E. cloacae patient isolate, ST191, blaKPC+ | (Chen et al. 2017) |
| ECNIH7 | E. cloacae patient isolate, ST53, blaKPC+ | (Chen et al. 2017) |
| xylR-L1 | K. pneumoniae, from evolution lineage 1, contains a missense SNP at xylR gene | This study |
| xylR-L2 | K. pneumoniae, from evolution lineage 2, contains a missense SNP at xylR gene | This study |
| xylR-L3 | K. pneumoniae, from evolution lineage 3, contains a missense SNP at xylR gene | This study |
| xylR-L4 | K. pneumoniae, from evolution lineage 4, contains a missense SNP at xylR gene | This study |
| xylR-L5 | K. pneumoniae, from evolution lineage 5, contains a missense SNP at xylR gene | This study |
| xylR-Tn1 | K. pneumoniae, loss-of-function mutant of xylR gene, from MKP103 transposon library #KP10497 | (Ramage et al. 2017) |
| xylR-Tn2 | K. pneumoniae, loss-of-function mutant of xylR gene, from MKP103 transposon library #KP10499 | (Ramage et al. 2017) |
